# Exploiting Low-Rank Structure in Genomic Relationship Matrices for Computationally Efficient Optimum Contribution Selection

**DOI:** 10.64898/2026.10.07.757325

**Authors:** Jon Ahlinder, Patrik Waldmann

## Abstract

**Motivation:** Optimum contribution selection (OCS) maximizes genetic gain under a constraint on relatedness. Genomic relationship matrices are dense, so the quadratic term costs *O*(*n*^3^) operations, and existing OCS software assumes pedigree sparsity.

**Results:** We replace the relationship matrix by its leading eigen-pairs, computed by randomized SVD, and solve OCS in factorized form. On Norway spruce population (*n* = 5,525), ranks 10–30 kept the exact objective within 1% of the optimum at 110– 330-fold speedups; on the QTL-MAS 2010 (*n* = 3,226), ranks up to 100 were needed (24–211-fold). The required rank rises as the diversity constraint loosens. Gain alone is a misleading accuracy check, since coancestry exceeded the optimum by up to 28%, so we choose the rank on the exact objective, which can be checked without a dense solve. The speedup comes from computing only the leading eigenpairs. We extend the factorization to robust OCS (18-fold).

**Availability and implementation:** Julia implementation, and examples are available at https://github.com/jonhar97/fastocs under the MIT license (archived at Zenodo, DOI 10.5281/zenodo.23214804).

**Supplementary information:** Supplementary data are available at *Bioinformatics* online.

## Introduction

Optimum contribution selection (OCS) (1) maximizes genetic gain while constraining the rate of inbreeding, and is a cornerstone of animal, plant and tree breeding. It is a quadratic program in which the contributions of selection candidates are optimized subject to a constraint on the expected mean kinship of the next generation (1, 2).

The shift from pedigree to genomic relationship matrices has improved the accuracy of genetic evaluation (3), but has removed the sparsity that made OCS tractable. Pedigree-based numerator relationship matrices (**A**) are sparse and admit exact recursive inversion, whereas genomic relationship matrices (**G**) are fully dense. The storage of **G** is *O*(*n*^2^) and standard matrix operations are *O*(*n*^3^) (4). Single-step **H** matrices, which blend pedigree and marker information for partially genotyped populations (5, 6), inherit this density.

Widely used OCS implementations, including GENCONT (7), optiSel (8), Opsel (9) and AlphaMate (10), were developed with sparse pedigree matrices in mind. Scalable reformulations built on that sparsity, such as block decomposition of **A** (11) and second-order cone programming on its sparse inverse (9) do not extend to a dense **G**, whose inverse is also dense. In practice, breeding programs pre-select or truncate the candidate list, sacrificing the premise of genomic informed OCS (that the whole genotyped candidate pool is evaluated), so that the optimal set of parents is found rather than an optimal subset of an arbitrarily reduced list.

Low-rank factorization addresses this directly. Relationship and covariance matrices arising in structured breeding populations are typically not truly of full rank. Limited effective population size, stratification and family structure concentrate their dominant structure in a small number of leading components (12, 13), which might result in a singular matrix. Retaining the top *k* singular triplets gives the best rank-*k* approximation in both the Frobenius and spectral norms (14). The same idea underpins factor-analytic covariance models in portfolio optimization (15, 16), whose mean–variance problem has the structure of OCS, so that it can be written as **G** *≈* **FF**^⊤^ + **D** which lets the quadratic form be evaluated as

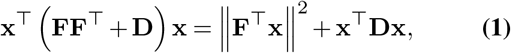

at *O*(*nk*) rather than *O*(*n*^2^) per evaluation, without materializing **G**. Computing only the leading *k* eigenpairs, by randomized SVD (RSVD hereafter) (17) or a Krylov (Lanczos) method, reduces the upfront cost from *O*(*n*^3^) to *O*(*n*^2^*k*). To our knowledge this combination has not been applied to OCS. We therefore (i) benchmark RSVD-OCS against the full-dense formulation on genetic gain, constraint satisfaction and solution concordance, (ii) separate the cost of the decomposition from that of the solve by comparing RSVD with a full eigendecomposition and with Lanczos, (iii) examine how the required rank depends on the tightness of the diversity constraint and how it can be chosen without a dense solve, and (iv) extend the factorization to robust OCS, which accounts for uncertainty in the estimated breeding values.

## Materials and Methods

### OCS and its low-rank reformulation

Let **c** ∈ℝ^*n*^ be the vector of contributions of *n* candidates to the next generation, **g** their estimated breeding values (EBVs) and **G** their relationship matrix. OCS solves

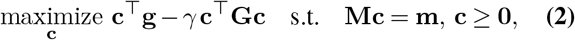

where **c**^⊤^**Gc** is twice the expected genomic coancestry of the next generation and *γ >* 0 trades gain against relatedness; it is chosen to correspond to a target rate of inbreeding (18). The linear constraints **Mc** = **m** are ∑_*i*_ *c*_*i*_ = 1 for the monoecious Norway spruce case and the two equalities ∑_*i∈S*_ *c*_*i*_ = ∑ _*i∈D*_*c*_*i*_ = 50 for the dioecious QTL-MAS case, where contributions are scaled to sum to 100, so that *γ* values are not comparable between the two datasets.

Because **G** is symmetric positive semidefinite, its SVD coincides with its eigendecomposition, and the rank-*k* truncation 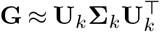 gives the factor 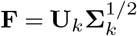. With the exact diagonal correction **D** = diag(**G**) *−* rowsums(**F** ◦**F**) ≥**0** and the auxiliary variable **y** = **F**^⊤^**c**, Eq. (1) turns Eq. (2) into a quadratic program in (**c, y**) *∈* ℝ^*n*^ *×*ℝ^*k*^ with objective **c**^⊤^**g***− γ*( ∥**y** ^2^∥ + **c**^⊤^**Dc**) and the additional linear equality **y** = **F**^⊤^**c** (Supplementary Note 2). Each evaluation of the quadratic term then costs *O*(*nk*) instead of *O*(*n*^2^), and the optimization stores *O*(*nk*) instead of *O*(*n*^2^) values. The decomposition costs *O*(*n*^3^) as a full eigendecomposition and *O*(*n*^2^*k*) by randomized SVD, where the latter accesses **G** only through products **GX**, which for a marker-based **G** = **ZZ**^⊤^*/s* can be formed as **Z**(**Z**^⊤^**X**)*/s* in *O*(*nmk*) for *m* markers without storing **G**.

We obtain **F** in three ways. First, *RSVD* (17) uses a Gaussian test matrix to sketch **G**, *q* = 2 rounds of subspace iteration with QR re-orthonormalisation refine an orthonormal basis **Q** (oversampling 10), and the small symmetric projection **B** = **Q**^⊤^**GQ** is eigendecomposed exactly (Supplementary Note 2). The two-sided projection is required for symmetric matrices; a one-sided sketch identifies the correct subspace but not the eigenvectors within it. Because the test matrix is random, RSVD has a stochastic component, which we quantify by replication with recorded seeds. Second, we used *Lanczos*, which computes the *k* leading eigenpairs with a deterministic Krylov method (KrylovKit.jl (19); tolerance 10^*−*8^). Third, in *Full eigen* the complete symmetric eigende-composition is used, but truncated to rank *k*.

### Rank selection

Every approximate solution **ĉ**_*k*_ was evaluated against the true **g** and **G**. Because Eq. (2) trades gain against coancestry, a solution can match the optimal gain while exceeding the optimal coancestry, so we measure accuracy by the exact objective gap

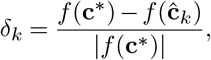

with *f* (**c**) = **c**^⊤^**g***− γ* **c**^⊤^**Gc** and **c**^*\**^ the full-dense solution, and report the gain and coancestry differences, the selection overlap *O*_*k*_ and the speedup *S*_*k*_ alongside it. The recommended rank is the smallest *k* with mean *δ*_*k*_ *<* 1% over replicates. Four alternative rules ((i) gain difference alone below 1%, (ii) *δ*_*k*_ *<* 0.5%, (iii) gain below 1% and coancestry within 5%, and (iv) the stabilisation rule below) are compared in Supplementary Note 2.

These rules need the full-dense solution and serve to validate the method. In routine use, *k* can be chosen without it, because **ĉ**_*k*_ is sparse, *f* (**ĉ**_*k*_) needs only the *s×s* submatrix of **G** for its *s* selected candidates, costing *O*(*s*^*2*^). We therefore also record the smallest tested *k* at which this exact objective changes by less than 0.1% to the next tested rank (stabilisation rule).

### Robust OCS. OCS treats **ĝ** as exact, ignoring its prediction error covariance **Ω** = Var(**g**|**y**). Following the robust reformulation of Fogg et al. (20) over an ellipsoidal uncertainty set,

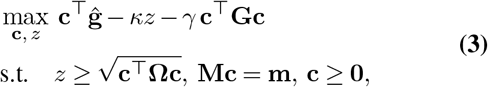

where *z* is an epigraph variable for the uncertainty term, *κ*≥0 is a robustness parameter and *κ* = 0 recovers standard OCS. Each sequential quadratic programming (SQP) step replaces the conic constraint by its tangent plane at the current iterate **c**_(*k*)_,

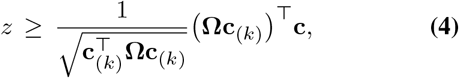

giving a convex QP with accumulated cuts. Because **G** and **Ω** enter Eq. (3) through separate terms, they are factorized independently, each with its own rank, which reduces the cost of computing the cut coefficients **Ωc**_(*k*)_ in Eq. (4) from *O*(*n*^2^) to *O*(*nr*_Ω_). Here **Ω** is the AI-REML prediction error covariance matrix, which is diagonally dominant rather than low rank and therefore takes the same low-rank-plus-exact-diagonal form used for **G**. Supplementary Note 2 gives the construction and contrasts it with the exactly low-rank factor available from an MCMC posterior sample (21).

### Data

*Norway spruce*. Two Skogforsk progeny trials in northern Sweden (S23F8820476 near Vindeln and S23F8820477 near Hädanberg), established in 1988 in un-replicated single-tree plots with offspring of 150 full-sib families, together with 820 unrelated, unimproved material. Of the *n* = 5,525 trees, 1,218 were genotyped by exome capture, and the single-step **H** matrix combining their genomic relationships with the pedigree was built as in Ahlinder and Waldmann (21), Chen et al. (22). Variance components, EBVs and their prediction error covariance matrix for tree height at age 17 (Hjd17; 4,462 trees phenotyped) were estimated by AI-REML (23), with trial as fixed effect and plot within trial, family within trial and the additive genetic effect (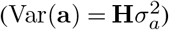) as random effects (Supplementary Note 1). The EBVs enter Eq. (2) as **g** and the prediction error covariance matrix enters Eq. (3) as **Ω**.

*QTL-MAS 2010*. The simulated pedigree of the 14th QTLMAS workshop (24), with *n* = 3,226 individuals over five generations, 9,723 SNPs after quality control, phenotypes for the first four generations (*n*_pheno_ = 2,326) and none for the fifth (*n*_pred_ = 900), with known true breeding values (TBV) and recorded sex. GEBVs for all pedigree members were estimated by GBLUP fitted with AI-REML, with sex as fixed effect (Supplementary Note 3), and all members were eligible candidates.

### Implementation and benchmarking

All models were implemented in Julia 1.11.4 (25) with JuMP 1.24.0 (26) and solved with OSQP (OSQP.jl 0.8.1) (27), a first-order ADMM solver that handles dense quadratic objectives and is robust to the mild ill-conditioning of near-singular relationship matrices (28). The full-dense baseline and the factorized models used identical solver settings (absolute and relative tolerance 10^*−*6^, at most 20,000 iterations, solution polishing, no warm start). To separate modelling-layer overhead from the solver, every model was also built and solved through the OSQP API directly, without JuMP. RSVD and Lanczos replicates used fixed, recorded seeds with *n* = 10 RSVD and *n* = 3 Lanczos and full-eigen replicates per rank in the Norway spruce single-*γ* comparison, and *n* = 5 RSVD replicates in the *γ ×*rank sweeps. The full-dense baseline was timed 2–3 times per *γ*. Wall-clock time was measured with Julia’s @elapsed, covering decomposition, model construction and solve, after warm-up solves on a small sub-problem that excluded compilation from all reported times. Benchmarks ran on a laptop with an Intel Core Ultra 7 155U processor (12 cores, 14 logical processors) and 16 GB RAM under Windows 11 (64-bit), with one Julia thread and 7 BLAS threads. Solution concordance was assessed by Spearman *ρ* and Kendall *τ* between contribution vectors, restricted to individuals selected (contribution *>* 10^*−* 4^) by both methods compared, since an unfiltered correlation is dominated by agreement on the unselected majority. Constraint satisfaction was assessed by evaluating the exact **c**^⊤^**Gc** at each approximate solution. Code is available at https://github.com/jonhar97/fastocs.

## Results

### Norway spruce

At the operating point *γ* = 400 the full-dense solution selected 73 of 5,525 candidates (1.3% with an expected coancestry of ^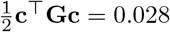^) in 331 s (SD 4.4, *n* = 3). The objective gap fell below 1% at rank 25 (0.98%), where RSVD-OCS was 136*×* faster (SD 14, *n* = 10) with 3.5% higher coancestry and 91.8% selection overlap. At rank 30 the gap was 0.10% and coancestry matched the optimum (Table 1, Figure 1). Gain alone would have suggested rank 10 (0.41%), but there coancestry exceeded the optimum by 28% and the objective gap was 6.0%: the truncated **G** under-represents relationships among the selected candidates, and the optimizer spends the slack on more related, higher-merit individuals. Speedups through the OSQP API directly were within 7% of those through JuMP.

**Table 1.**
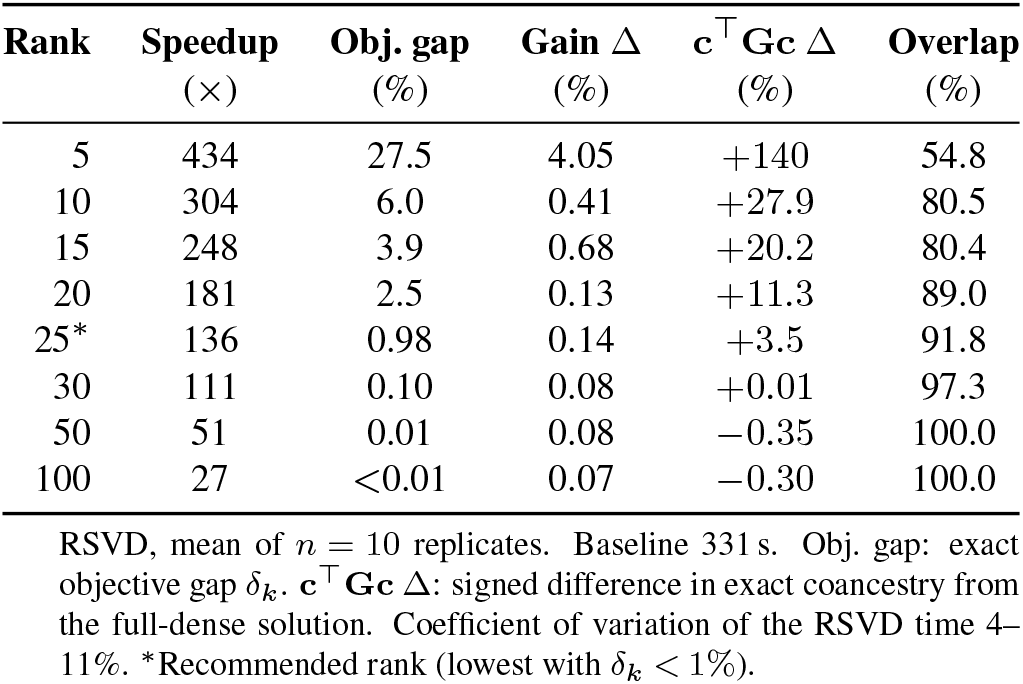
Rank comparison, Norway spruce (*γ* = 400)

| Rank | Speedup<br>(×) | Obj. gap<br>(%) | Gain $\Delta$<br>(%) | $c^T G c \Delta$<br>(%) | Overlap<br>(%) |
| --- | --- | --- | --- | --- | --- |
| 5 | 434 | 27.5 | 4.05 | +140 | 54.8 |
| 10 | 304 | 6.0 | 0.41 | +27.9 | 80.5 |
| 15 | 248 | 3.9 | 0.68 | +20.2 | 80.4 |
| 20 | 181 | 2.5 | 0.13 | +11.3 | 89.0 |
| 25* | 136 | 0.98 | 0.14 | +3.5 | 91.8 |
| 30 | 111 | 0.10 | 0.08 | +0.01 | 97.3 |
| 50 | 51 | 0.01 | 0.08 | -0.35 | 100.0 |
| 100 | 27 | <0.01 | 0.07 | -0.30 | 100.0 |

RSVD, mean of $n = 10$ replicates. Baseline 331 s. Obj. gap: exact objective gap $\delta_k$ . $c^T G c \Delta$ : signed difference in exact coancestry from the full-dense solution. Coefficient of variation of the RSVD time 4–11%. \*Recommended rank (lowest with $\delta_k < 1\%$ ).

**Fig. 1.**
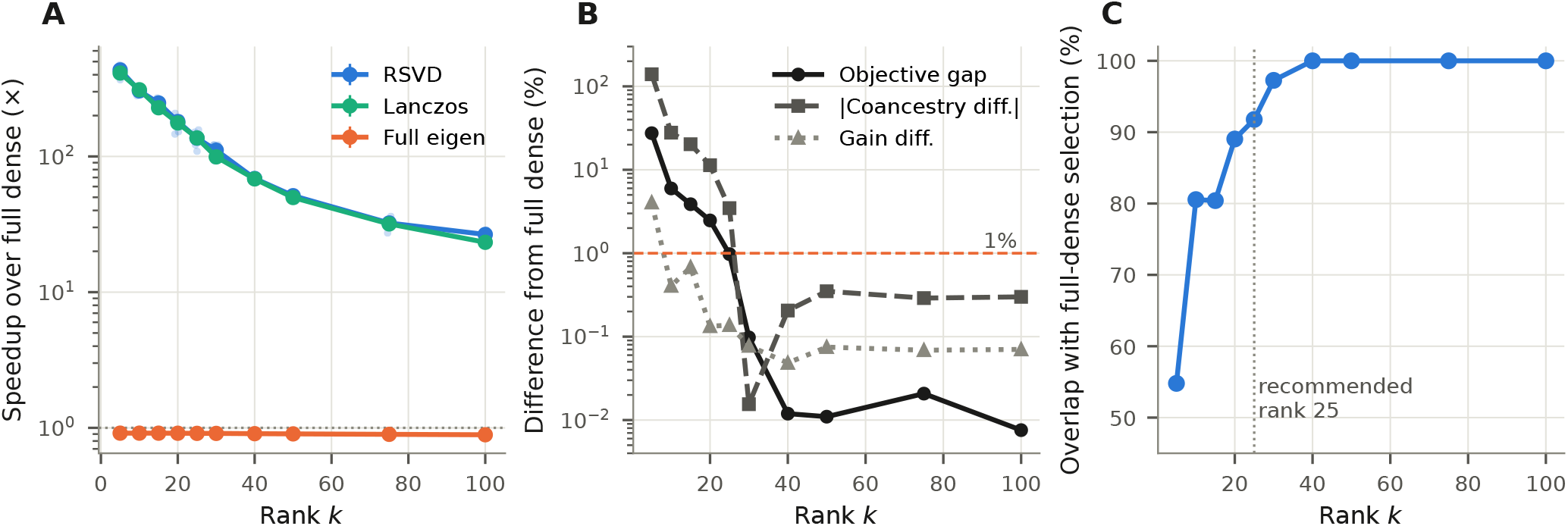
Rank selection, Norway spruce, *γ* = 400. **(A)** Speedup over the full-dense baseline (331 s) for RSVD (*n* = 10 replicates, translucent points), Lanczos and the full eigendecomposition (*n* = 3 each) with means *±* SD. The dotted line marks equal time. **(B)** Exact objective gap, absolute coancestry difference and gain difference of the RSVD solutions relative to the full-dense solution (log scale, with values below 0.005% shown at 0.005%). The dashed line marks 1%. **(C)** Overlap between the RSVD and full-dense selected sets.

Lanczos matched RSVD in accuracy and speed (137*×* at rank 25), whereas the full eigendecomposition took 361 s and made the method slower than the dense solve (0.91*×*), although its factorized solve was as fast. The speedup therefore comes from computing only the leading *k* eigenpairs, not from randomization, where RSVD kept a small advantage at high rank (1.2 versus 3.0 s at rank 100). At rank 30 the jointly selected individuals were ranked almost identically by the approximation and the baseline (Spearman *ρ* = 0.967, *n* = 71 Supplementary Figure S1, Table S1). The exact objective of the approximate solutions, computed from the selected candidates alone, changed by less than 0.1% from rank 30 onwards, so the stabilisation rule selected rank 30 without a dense solve.

Across *γ*, the rank needed rose from 10 to 30 as the diversity constraint loosened and the full-dense solution grew from 20 to 348 candidates, and the speedup at that rank (110–331*×*) declined with the number selected (log–log *r* = − 0.84; Figure 2, Table 2). The gain-only rule would have left objective gaps of 0.9–6.1%, whereas the stabilisation rule never chose a lower rank than the 1% objective rule (Supplementary Note 2). Above rank 40 the gap levelled off at 0.1–0.25% for *γ*≥ 1000, the resolution limit of the baseline’s solver tolerance. The sweep, run in a separate session, gave 233*×* at *γ* = 400 and rank 25 against 136*×* above. Within a session, the timing varied by 4–11%.

**Table 2.** Recommended rank per diversity constraint strength, Norway spruce (*n* = 5,525) and QTL-MAS 2010 (*n* = 3,226)

| Dataset | $\gamma$ | Sel. | Rank | Speedup (×) | Obj. gap (%) | $\mathbf{c}^\top \mathbf{G} \mathbf{c} \Delta$ (%) | Overlap (%) |
| --- | --- | --- | --- | --- | --- | --- | --- |
| Norway spruce | 100 | 20 | 10 | 331 (SD 182) | 0.63 | +3.0 | 90.0 |
|  | 200 | 35 | 15 | 223 (SD 15) | 0.90 | +7.2 | 82.9 |
|  | 400 | 73 | 25 | 233 (SD 43) | 0.98 | +3.4 | 91.8 |
|  | 1000 | 148 | 30 | 243 (SD 21) | 0.32 | +0.6 | 95.3 |
|  | 3000 | 348 | 30 | 110 (SD 21) | 0.90 | +1.7 | 95.7 |
| QTL-MAS 2010 | 0.1 | 13 | 5 | 211 (SD 52) | 0.63 | +7.0 | 92.3 |
|  | 0.5 | 43 | 50 | 56 (SD 4) | 0.84 | +2.5 | 90.7 |
|  | 1 | 78 | 100 | 24 (SD 3) | 0.96 | +0.3 | 89.5 |
|  | 2 | 145 | >100 | 27 (SD 9) | 3.1 | +7.5 | 76.0 |
|  | 5 | 325 | >100 | 37 (SD 1) | 14.8 | +35 | 66.0 |
|  | 10 | 567 | >100 | 33 (SD 1) | 56.0 | +118 | 54.2 |

Sel.: individuals with contribution $> 10^{-4}$ in the full-dense solution. Rank: lowest rank with a mean exact objective gap $< 1\%$ over five RSVD replicates ( $> 100$ ): not reached within the tested ranks, values shown at rank 100. QTL-MAS contributions sum to 100, so $\gamma$ is not comparable between datasets. Full-dense baseline 188–335 s (Norway spruce) and 26–31 s (QTL-MAS) through JuMP. Overlap: % of full-dense-selected individuals also selected by RSVD.

**Fig. 2.**
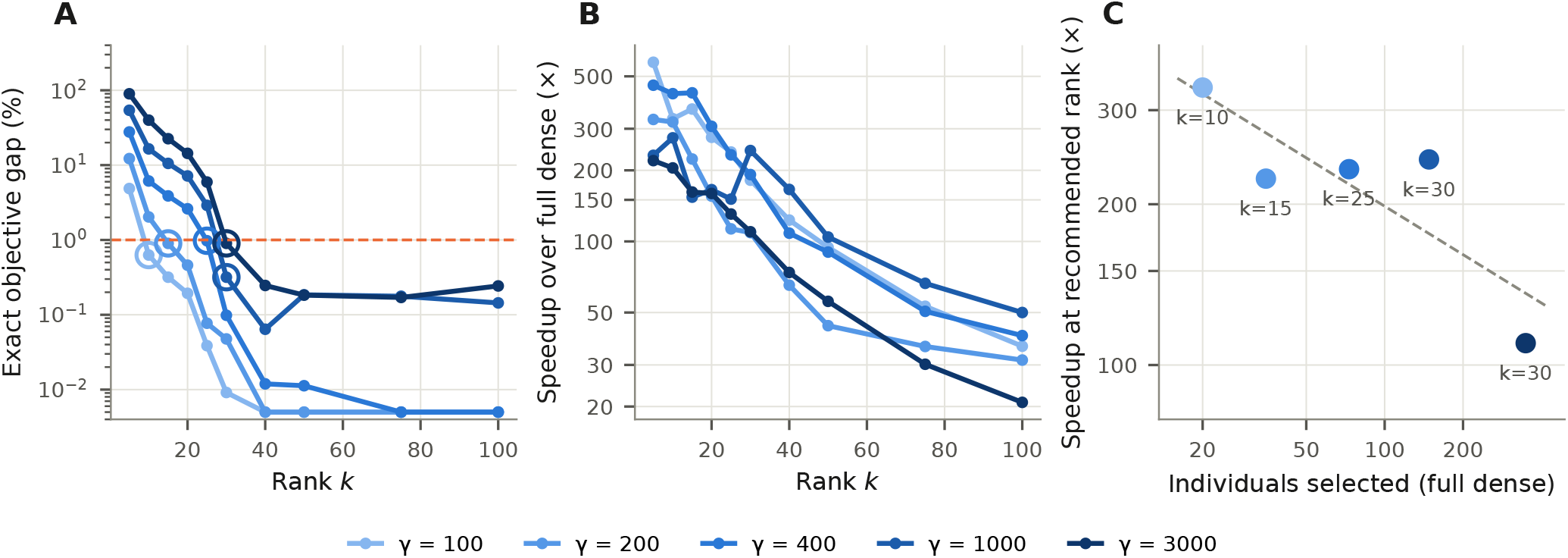
Effect of diversity constraint strength on the accuracy and speedup of RSVD-OCS (Norway spruce, *n* = 5,525). **(A)** Exact objective gap between RSVD-OCS and the full-dense baseline against rank *k* for five values of *γ* (mean of five replicates; log scale, values below 0.005% shown at 0.005%). The dashed line marks 1%, and open circles mark the recommended rank per *γ*. **(B)** Speedup over the full-dense solver against rank. **(C)** Speedup at the recommended rank against the number of individuals selected in the full-dense solution, labelled with the recommended rank; the dashed line is a log–log fit.

### Robust OCS

At the same operating point (*γ* = 400, *r*_*G*_ = 30), the AI-REML prediction error covariance **Ω** was not low rank (its 100 largest eigenvalues hold 10% of its trace), but the low-rank-plus-diagonal form at *r*_Ω_ = 30 reproduced the exact uncertainty term to within 0.6%, and all values in Table 3 are exact.

**Table 3.**
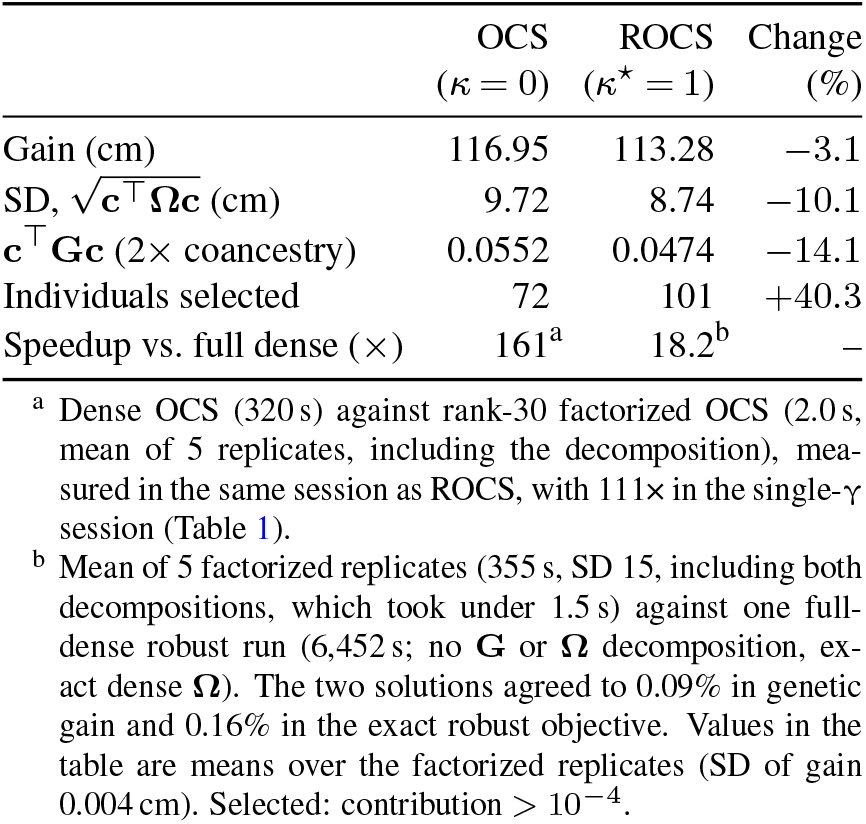
Robust OCS results, Norway spruce (*γ* = 400, *r*_*G*_ = 30, *r*_Ω_ = 30 plus exact diagonal correction)

| | OCS<br>($\kappa = 0$) | ROCS<br>($\kappa^* = 1$) | Change<br>(%) |
| --- | --- | --- | --- |
| Gain (cm) | 116.95 | 113.28 | −3.1 |
| SD, $\sqrt{\mathbf{c}^\top \mathbf{\Omega} \mathbf{c}}$ (cm) | 9.72 | 8.74 | −10.1 |
| $\mathbf{c}^\top \mathbf{G} \mathbf{c}$ ($2 \times$ coancestry) | 0.0552 | 0.0474 | −14.1 |
| Individuals selected | 72 | 101 | +40.3 |
| Speedup vs. full dense (×) | 161<sup>a</sup> | 18.2<sup>b</sup> | — |

<sup>a</sup> Dense OCS (320 s) against rank-30 factorized OCS (2.0 s, mean of 5 replicates, including the decomposition), measured in the same session as ROCS, with $111\times$ in the single- $\gamma$ session (Table 1).
<sup>b</sup> Mean of 5 factorized replicates (355 s, SD 15, including both decompositions, which took under 1.5 s) against one full-dense robust run (6,452 s; no $\mathbf{G}$ or $\mathbf{\Omega}$ decomposition, exact dense $\mathbf{\Omega}$ ). The two solutions agreed to 0.09% in genetic gain and 0.16% in the exact robust objective. Values in the table are means over the factorized replicates (SD of gain 0.004 cm). Selected: contribution $> 10^{-4}$ .

ROCS traded a 3.1% loss in gain for 10.1% lower uncertainty and 14.1% lower coancestry, spreading contributions over 101 rather than 72 individuals. Its speedup (18.2*×*) was well below the 161*×* of standard OCS in the same session, where each SQP iteration took about 23 s against 1.5 s for a single factorized OCS solve, because every cut (Eq. (4)) is a dense length-*n* constraint row and the QP is rebuilt with all accumulated cuts at each iteration. Neither factorization removes this cost. Both SQPs converged to the solver tolerance within 14–15 iterations.

### QTL-MAS 2010

GBLUP by AI-REML gave *ĥ*^*2*^= 0.446 (95% CI 0.386–0.507), consistent with the simulated heritabilities (24), and GEBV accuracies of *r* = 0.702 (phenotyped) and 0.607 (unphenotyped) (Supplementary Note 3). The rank needed again rose as the constraint loosened, but more steeply than for Norway spruce: from 5 at *γ* = 0.1 (211 *×*) to 100 at *γ* = 1 (24 *×*), with no tested rank sufficient for *γ≥* 2 (Table 2, Supplementary Figure S3). At the operating point *γ* = 1 (78 selected, Δ*F ≈*2.9% per generation), rank 30 passed on gain (0.42%) but exceeded the optimal coancestry by 19%, where the stabilisation rule did not settle within rank 100, correctly signalling that a higher rank was needed.

At rank 100, 72 of the 78 individuals selected by the full-dense solution were also selected by RSVD (Spearman *ρ* = 0.888 among them, Supplementary Figure S4, Table S3). Because TBV are known, we checked whether disagreements matter, and individuals selected by only one method did not differ in TBV from those selected by both (*p* = 0.23 and *p >* 0.99), all far above the unselected majority (Supplementary Figure S5), and the contribution-weighted TBV differed by 0.3% (28.71 versus 28.61).

## Discussion

Genomic relationship matrices have replaced pedigree matrices as the standard for inference in animal models, but this transition exposed dense matrix computation as a barrier that existing OCS software was never designed to handle. Replacing **G** by its leading eigenpairs within the OCS quadratic program gave speedups of one to two orders of magnitude on two case studies differing in size and structure, with the exact objective within 1% of the optimum. Two results qualify how the method should be used. First, the required rank depends on the data and on the diversity constraint, with 10– 30 sufficed for Norway spruce, but QTL-MAS needed up to 100 at comparable selection intensity and more under weak constraints. Second, genetic gain is not a sufficient accuracy check, because a truncated **G** understates relationships among the selected candidates and lets the optimizer trade diversity for gain. Thus, the exact objective, or gain together with coancestry, must be checked. The stabilisation rule does this without a dense solve and was conservative on both datasets.

The speedup does not depend on randomization. Lanczos matched RSVD in accuracy and nearly in speed, and a full eigendecomposition was slower than the dense solve. What matters is that only the leading *k* eigenpairs are computed. RSVD is attractive because it needs a fixed, small number of passes over **G**, which suits out-of-core and distributed settings (17), but a Krylov method, as used for biobank-scale genotype data (29), is an equally valid choice.

The practical relevance is best seen against current population sizes. Genotyped candidate pools of several hundred thousand individuals are routine in cattle breeding (30), and tree breeding is moving the same way (31, 32) as genotyping costs fall. At that scale dense OCS is not merely slow but infeasible, and programs respond either by truncating the candidate list or by substituting the sparse **A** matrix for **G**. Both sacrifice the premise of genomic OCS. Because a moderate rank preserved the objective and selection membership on both datasets, the entire genotyped candidate pool can be retained in the optimization, provided the rank is checked as above. At that scale **G** itself can no longer be stored, but RSVD needs only products with **G**, which for a marker-based **G** can be formed from the marker matrix; a single-step **H** still requires its dense genotyped block. Furthermore, we did not test RSVD-OCS on populations larger than 5,525, where the case for larger pools rests on the *O*(*nk*) memory and *O*(*n*^2^*k*) decomposition costs (17), and the rank needed at that scale will depend on the population’s structure (33).

Three limitations bound these conclusions. We tested two populations, one trait per population and one construction of **G** each, and the attainable speedup depends on the effective rank of **G** and on how sparse the optimal contribution vector is under the chosen diversity constraint; other constructions of the relationship matrix, for which no consensus exists in genomic OCS (34), may need different ranks. Timings came from one laptop and varied by up to 1.6-fold between sessions, so the speedups are order-of-magnitude figures. And accuracy was measured against a first-order solver at tolerance 10^*−*6^, which limits the resolvable objective gap to about 0.1–0.2%.

Our use of low-rank approximation follows a broader trend in genomic evaluation. Meyer (4, 12) showed that reparameterizing the mixed model equations onto a principal-component scale reduces REML iteration cost, resting on the same dimensionality argument as our rank selection, and related reformulations avoid dense inversion by re-expressing or accelerating the linear algebra of variance component estimation (30, 35). Our contribution applies the principle inside the optimization itself rather than in the REML/BLUP step that produces its input, so the approaches are complementary, where a rank-reduced **G** could be paired with a PC-reparameterized mixed model to compress cost across the whole pipeline. Among alternatives that approximate **G** directly, the most widely adopted is APY (36), which partitions genotyped individuals into an exactly-inverted core and a conditionally independent remainder, with core size scaling with the effective number of independent chromosome segments (33). APY is a deterministic, individual-selection approximation built to provide a sparse inverse for genomic evaluation, whereas RSVD uses a global random projection, requires no such selection and targets the evaluation of the OCS objective rather than the inverse; we therefore do not compare the two directly. A contemporaneous application of randomized PCA to large pedigree matrices (37) suggests randomized approximation is gaining traction in this literature more broadly.

For the robust extension, Fogg et al. (20) give a formulation that is exact with respect to **Ω** but, like classical OCS with respect to **G**, does not address its *O*(*n*^2^) cost. The same factorization principle extends there, with each matrix exploiting whatever structure its own source provides. The speedups do not multiply, however: each SQP iteration adds a dense length-*n* cut and rebuilds the QP, a cost that neither factorization removes and that becomes binding once **G**’s contribution is compressed. The contribution here is computational, and the modelling of breeding-value uncertainty itself is addressed elsewhere (20, 21). The general lesson is that the constraint structure an extension introduces, not only which matrices are approximated, determines how much of the lowrank speedup carries through.

A separate lineage has scaled the optimization step rather than the matrix: exploiting pedigree block structure (11), replacing the Lagrangian solve with Newton-Raphson iteration (38), or recasting OCS as semidefinite (39, 40) or second-order cone programming (9), binary integer programming or convex mate allocation (42). RSVD-OCS is orthogonal to these, so that it changes what is fed into the solver rather than how the problem is solved, and could be combined with any of them.

## Conclusions

The repeated dense evaluation of **c**^⊤^**Gc**, rather than any single operation, is what makes genomic OCS costly as breeding populations scale. Approximating the relationship matrix by its leading eigenpairs removes that cost. On the analyzed Norway spruce population, ranks 10–30 kept the exact objective within 1% of the optimum while cutting computation time 110–330-fold, and on QTL-MAS ranks up to 100 were needed. The required rank grows as the diversity constraint loosens, must be judged on the exact objective rather than on gain alone, and can be checked without a dense solve. The same factorization extends to robust OCS, where breeding-value uncertainty enters as a second, independently factorized covariance matrix, showing that the principle generalises beyond the relationship matrix to other structured matrices in the OCS objective. As genotyped candidate pools grow, low-rank approximation offers a general way to keep OCS tractable without sacrificing its optimality guarantee, and pairs naturally with the solver-side and trait-space reductions emerging elsewhere (34, 43).

## Author contributions

Jon Ahlinder (Conceptualization [equal], Formal analysis [lead], Software [lead], Writing – original draft [lead]), Patrik Waldmann (Conceptualization [equal], Software [supporting], Supervision [lead], Writing – review & editing [lead]).

## Conflict of interest

None declared.

## Funding

J.A. was supported by project ID 21414 (Föreningen ID 23-511), funded by Föreningen Skogsträdsförädling.

## Notes

https://doi.org/10.5281/zenodo.23214804

https://github.com/jonhar97/fastocs

https://doi.org/10.5281/zenodo.23206385

### Competing Interest Statement

The authors have declared no competing interest.

