## Supplementary material for "Exploiting Low-Rank Structure in Genomic Relationship Matrices for Computationally Efficient Optimum Contribution Selection"

DRAFT

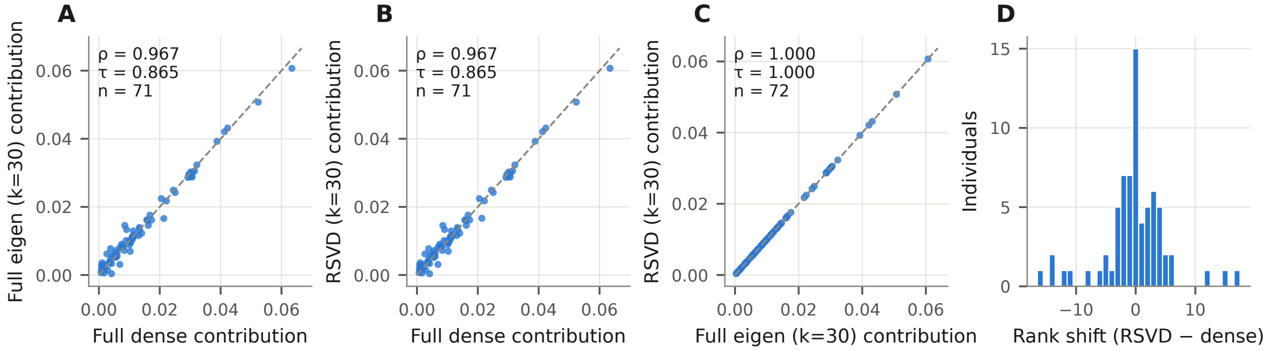


**Fig. S1. Solution concordance between approximation methods and the full-dense baseline, Norway spruce (rank 30, $\gamma = 400$).** All panels restricted to individuals selected (contribution $> 10^{-4}$) by both methods being compared, avoiding inflation from the majority of the pedigree both methods agree is unselected. **(A)** Truncated full eigendecomposition vs. full dense ($\rho = 0.967$, $\tau = 0.865$, $n = 71$). **(B)** RSVD (replicate 1) vs. full dense ($\rho = 0.967$, $\tau = 0.865$, $n = 71$). **(C)** RSVD vs. truncated full eigendecomposition ($\rho = 1.000$, $\tau = 1.000$, $n = 72$). **(D)** Rank shifts (RSVD − full dense) among the 71 jointly selected individuals (mean absolute shift 3.5; maximum 17).

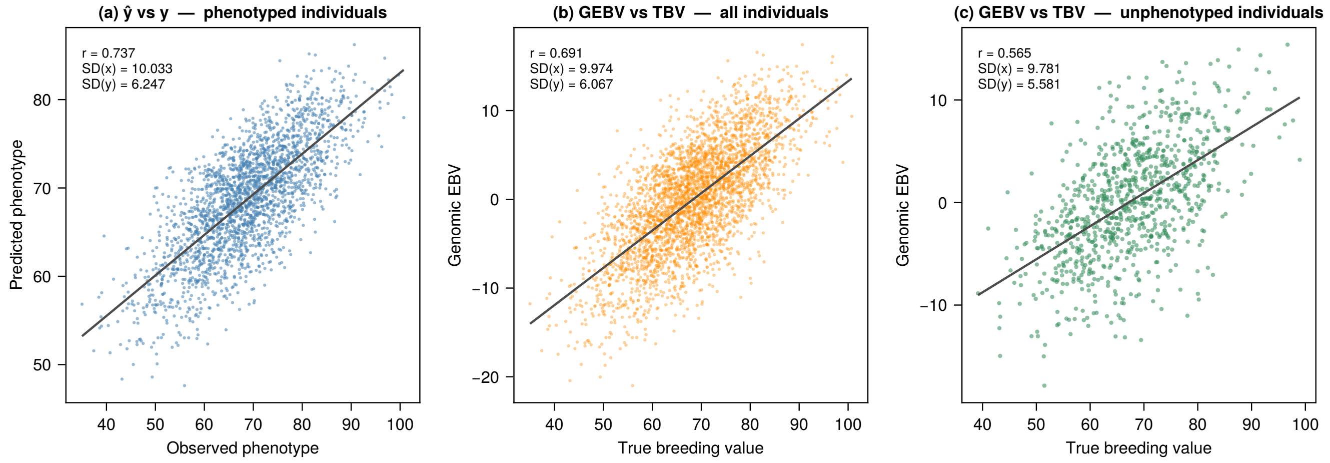


**Fig. S2.** GBLUP prediction accuracy for the QTL-MAS 2010 simulated dataset. (a) Predicted versus observed phenotype for the $n_{\text{pheno}} = 2{,}326$ training individuals. (b) GEBV versus TBV for all $n = 3{,}226$ pedigree members. (c) GEBV versus TBV for the $n_{\text{pred}} = 900$ unphenotyped fifth-generation individuals. Pearson $r$, SD of each axis, and OLS regression line shown in each panel.

Dr

**Table S1.** Solution quality metrics at rank 30, Norway spruce ($\gamma = 400$)

| **Metric** | **Full eigen** | **RSVD** | **Lanczos** |
|---|---|---|---|
| Spearman $\rho$ (jointly selected) | 0.967 | 0.967 | 0.967 |
| Kendall $\tau$ (jointly selected) | 0.865 | 0.865 | 0.865 |
| Spearman $\rho$ (unfiltered, $n = 5{,}525$) | 0.970 | 0.970 | 0.970 |
| Selection overlap with full dense (%) | 97.3 | 97.3 | 97.3 |
| Exact objective gap (%) | 0.098 | 0.098 | 0.098 |
| Genetic gain difference (%) | 0.08 | 0.08 | 0.08 |
| Coancestry difference (%) | +0.01 | +0.01 | +0.01 |
| Decomposition time (s) | 361 | 0.45 | 0.73 |
| Speedup (×) | 0.91 | 111 | 99 |
| Mean / max rank shift, jointly selected | – | 3.5 / 17 | – |

Jointly selected: contribution $> 10^{-4}$ under both the full-dense baseline and the method compared ($n = 71$). Concordance for RSVD from replicate 1; accuracy and time are means over replicates ($n = 10$ RSVD, $n = 3$ otherwise). Rank shifts for RSVD only (Figure S1D).

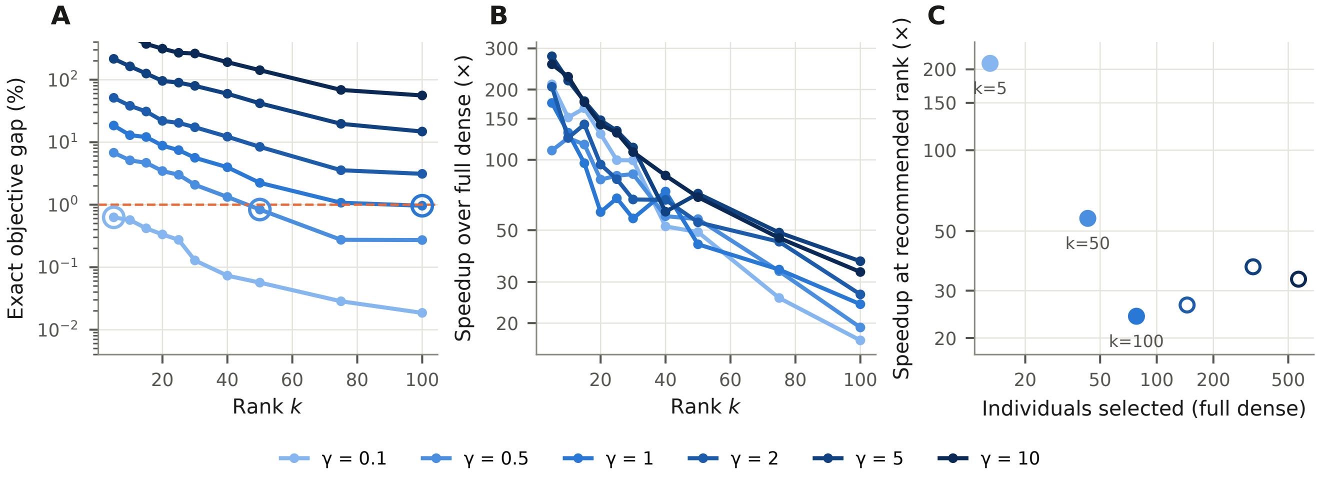


**Fig. S3. Effect of diversity constraint strength on the accuracy and speedup of RSVD-OCS (QTL-MAS 2010, $n = 3{,}226$). (A)** Exact objective gap between RSVD-OCS and the full-dense baseline against rank $k$ for six values of $\gamma$ (mean of five replicates, log scale). The dashed line marks 1%; open circles mark the recommended rank. Curves for $\gamma = 2$, 5 and 10 did not reach 1% within $k \leq 100$. **(B)** Speedup over the full-dense solver against rank; it remains above 17× in all conditions. **(C)** Speedup at the recommended rank against the number of individuals selected in the full-dense solution. Open points: no rank $\leq 100$ reached a gap below 1% (shown at $k = 100$). Tighter diversity constraints give sparser solutions, which are approximated well at lower rank and thus afford greater speedup.

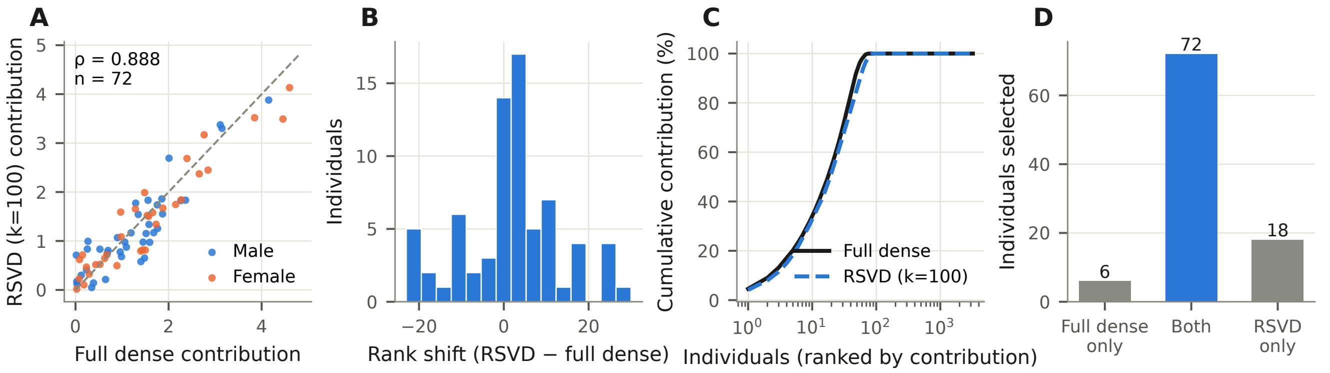


**Fig. S4. Solution concordance between RSVD-OCS (recommended rank 100, replicate 1) and the full-dense baseline for the QTL-MAS 2010 dataset ($\gamma = 1$). (A)** Contributions of the 72 individuals selected by both methods (Spearman $\rho = 0.888$), coloured by sex. **(B)** Rank shifts (RSVD − full dense) among the same individuals (mean absolute shift 8.9 positions; maximum 30). **(C)** Cumulative contribution across all $n = 3{,}226$ pedigree members. **(D)** Individuals selected by the full-dense solution only (6), by both (72) and by RSVD only (18).

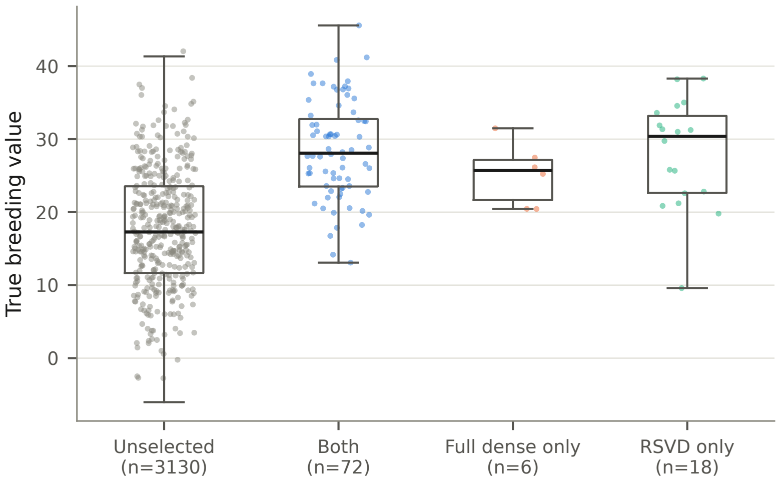


**Fig. S5. True breeding value (TBV) of selected versus unselected individuals, QTL-MAS 2010 ($\gamma = 1$, rank 100).** Individuals selected by both methods have substantially higher TBV than the unselected majority (Mann–Whitney $p = 6.6 \times 10^{-23}$). Individuals selected by only one method do not differ from those selected by both (full dense only: $p = 0.23$; RSVD only: $p > 0.99$). Points are a random subsample (at most 400 per group), jittered; boxes show median and interquartile range.

**Table S2.** GBLUP variance component estimates for the QTL-MAS 2010 dataset (AI-REML)

| Parameter | Estimate (95% CI) |
|---|---|
| $\sigma_G^2$ | 43.94 (34.29–53.59) |
| $\sigma_e^2$ | 54.51 (50.92–58.10) |
| $h^2$ | 0.446 (0.386–0.507) |
| Log-likelihood | $-6078.49$ |
| AIC / BIC | 12164.98 / 12187.99 |
| Accuracy, phenotyped | 0.702 |
| Accuracy, unphenotyped | 0.607 |

AI-REML converged in 6 iterations. Fixed effects: intercept $\hat{\beta} = 68.61$ (SE 0.245), gender $\hat{\beta} = 0.179$ (SE 0.326). Accuracy: Pearson correlation between GEBV and true breeding value, for the $n = 2{,}326$ phenotyped and $n = 900$ unphenotyped individuals.

**Table S3.** Solution quality metrics for the QTL-MAS 2010 dataset at rank 100, $\gamma = 1$

| Metric | RSVD | Full dense |
|---|---|---|
| Genetic gain ($\mathbf{c}^\top\mathbf{g}$) | 1214.2 | 1220.9 |
| $\mathbf{c}^\top\mathbf{G}\mathbf{c}$ | 289.3 | 288.7 |
| Objective $\mathbf{c}^\top\mathbf{g} - \gamma\mathbf{c}^\top\mathbf{G}\mathbf{c}$ | 924.9 | 932.3 |
| Selected (total) | 90 | 78 |
| Selected males / females | 50 / 40 | 41 / 37 |
| Jointly selected (both methods) | 72 | |
| Spearman $\rho$ (jointly selected, $n = 72$) | 0.888 | |
| Objective gap / gain difference (%) | 0.78 / $-0.55$ | – |
| Realised gain, TBV-weighted | 28.61 | 28.71 |

RSVD replicate 1 (mean over five replicates: objective gap 0.96%, gain difference 0.68%, 24× speedup). Contributions sum to 100 (50 per sex). Concordance restricted to individuals selected by both methods (Figure S4).

### Supplementary Note 1: Single-step HBLUP AI-REML analysis of Norway spruce height (Hjd17)

Breeding values used in the Norway spruce OCS analyses were obtained from a single-trait single-step HBLUP model for tree height at age 17 (Hjd17; $n = 4{,}462$ phenotyped of $5{,}525$ trees),

$$\mathbf{y} = \mathbf{X}\mathbf{b} + \mathbf{Z}_p\mathbf{p} + \mathbf{Z}_f\mathbf{f} + \mathbf{Z}_a\mathbf{a} + \mathbf{e},$$

with trial as fixed effect ($\mathbf{b}$), random plot-within-trial ($\mathbf{p} \sim N(\mathbf{0}, \mathbf{I}\sigma_p^2)$, 92 levels) and family-within-trial effects ($\mathbf{f} \sim N(\mathbf{0}, \mathbf{I}\sigma_f^2)$, 261 levels) capturing genotype-by-environment interaction, additive genetic effects $\mathbf{a} \sim N(\mathbf{0}, \mathbf{H}\sigma_a^2)$ and residuals $\mathbf{e} \sim N(\mathbf{0}, \mathbf{I}\sigma_e^2)$. Variance components were estimated by AI-REML, and EBVs and the full prediction error covariance matrix, $\mathrm{PEV} = \sigma_a^2\mathbf{H} - \sigma_a^4\mathbf{H}_{\cdot o}\mathbf{P}\mathbf{H}_{o\cdot}$ (where $o$ indexes phenotyped trees and $\mathbf{P}$ is the REML projection matrix), were computed for all $5{,}525$ trees. For comparison, the same model was fitted with the pedigree relationship matrix $\mathbf{A}$ (ABLUP) and with $\mathbf{H}$ but without the family-within-trial term. The data include 820 unrelated reference trees (646 phenotyped) of unimproved material, with genetic groups rather than individual trees recorded as parents; these enter $\mathbf{H}$ as unrelated base-population individuals.
The HBLUP model converged in 12 iterations and fitted the data better than ABLUP (Table S4). Omitting the family-within-trial term reduced the REML log-likelihood by 21.9 units (likelihood ratio statistic 43.8, 1 df, $p < 0.001$) and inflated $\hat{\sigma}_a^2$ by 24%. HBLUP EBVs were highly correlated with ABLUP EBVs and with the posterior means from the earlier multi-trait JWAS analysis (Table S5, Figure S6B,C), but were more strongly shrunk than the JWAS estimates (regression slope 0.65), which did not include the family-within-trial term. In five-fold cross-validation (random folds, variance components fixed at their full-data estimates), the predictive ability, i.e. the correlation between predicted genetic values and phenotypes adjusted for trial and plot effects, was 0.249 for HBLUP and 0.242 for ABLUP (0.222 and 0.227 excluding the reference trees), with a regression slope of 1.04 for HBLUP, indicating little bias in dispersion (Figure S6A).

### Supplementary Note 2: Low-rank reformulation, randomized SVD and rank selection

**Low-rank reformulation of the OCS quadratic program.** The genomic relationship matrix $\mathbf{G} \in \mathbb{R}^{n\times n}$ is dense and symmetric positive semidefinite, so its singular value decomposition coincides with its eigendecomposition and the rank-$k$ truncation is

$$\mathbf{G} \approx \mathbf{G}_k = \mathbf{U}_k\mathbf{\Sigma}_k\mathbf{U}_k^\top, \qquad \mathbf{F} = \mathbf{U}_k\mathbf{\Sigma}_k^{1/2} \in \mathbb{R}^{n\times k},$$

DRAFT

**Table S4. Variance components (SE) for Hjd17, Norway spruce.** REML log-likelihoods are comparable across models (same data and fixed effects).

| | HBLUP | HBLUP, no fam.×trial | ABLUP |
|---|---|---|---|
| $\sigma_a^2$ | 4396 (948) | 5438 (979) | 2748 (793) |
| $\sigma_p^2$ (plot within trial) | 2491 (422) | 2434 (410) | 2470 (419) |
| $\sigma_f^2$ (family within trial) | 625 (147) | – | 699 (161) |
| $\sigma_e^2$ | 9354 (650) | 9128 (664) | 10672 (521) |
| $h^2$ | 0.261 (0.051) | 0.320 (0.051) | 0.166 (0.045) |
| REML log-likelihood | −23432.4 | −23454.3 | −23440.5 |
| Mean reliability, phenotyped | 0.46 | – | 0.43 |
| Mean reliability, unphenotyped | 0.28 | – | 0.34 |

**Table S5. Agreement between EBVs for Hjd17 from HBLUP (AI-REML), ABLUP (AI-REML) and multi-trait JWAS.** Pearson $r$ / Spearman $\rho$; top-$k$: number of the $k$ highest-ranked trees shared.

| Comparison | All ($n = 5{,}525$) | Phenotyped (4,462) | Unphenotyped (1,063) | Top 100 | Top 200 |
|---|---|---|---|---|---|
| HBLUP vs. JWAS | 0.940 / 0.939 | 0.948 / 0.947 | 0.901 / 0.907 | 64 | 133 |
| HBLUP vs. ABLUP | 0.947 / 0.949 | 0.945 / 0.945 | 0.976 / 0.952 | 72 | 138 |
| ABLUP vs. JWAS | 0.848 / 0.857 | 0.853 / 0.853 | 0.889 / 0.903 | – | – |

with $\mathbf{U}_k$ the $k$ leading eigenvectors and $\mathbf{\Sigma}_k$ the corresponding eigenvalues. Writing $\mathbf{G} \approx \mathbf{F}\mathbf{F}^\top + \mathbf{D}$, where the diagonal correction $\mathbf{D} = \mathrm{diag}(\mathbf{G}) - \mathrm{rowsums}(\mathbf{F} \circ \mathbf{F})$ restores the exact self-relationships, and introducing $\mathbf{y} = \mathbf{F}^\top\mathbf{c} \in \mathbb{R}^k$, the quadratic term becomes $\mathbf{c}^\top\mathbf{G}\mathbf{c} \approx \|\mathbf{y}\|^2 + \mathbf{c}^\top\mathbf{D}\mathbf{c}$. The OCS problem is then a quadratic program in $(\mathbf{c}, \mathbf{y})$ with the original linear constraints plus the equality $\mathbf{y} = \mathbf{F}^\top\mathbf{c}$. Memory drops from $O(n^2)$ to $O(nk)$ and each evaluation of the quadratic form from $O(n^2)$ to $O(nk)$; $\mathbf{G}$ itself is never passed to the solver.

**Randomized SVD.** Computing the full decomposition of $\mathbf{G}$ costs $O(n^3)$. Randomized SVD (1) instead identifies a low-dimensional subspace capturing the dominant action of $\mathbf{G}$ and decomposes within it:

1. Draw $\mathbf{R} \in \mathbb{R}^{n\times\ell}$ with $R_{ij} \sim \mathcal{N}(0,1)$, where $\ell = k + p$ and $p = 10$ is the oversampling.
2. Form the sketch $\mathbf{Y} = \mathbf{G}\mathbf{R}$ and its orthonormal basis $\mathbf{Q}$ by QR decomposition.
3. Refine $\mathbf{Q}$ by $q$ rounds of subspace iteration, each $\mathbf{Q} \leftarrow \mathrm{qr}(\mathbf{G}\mathbf{Q})$ applied twice, with re-orthonormalisation after every product to avoid loss of precision (we use $q = 2$).
4. Form the small symmetric matrix $\mathbf{B} = \mathbf{Q}^\top\mathbf{G}\mathbf{Q} \in \mathbb{R}^{\ell\times\ell}$ and eigendecompose it exactly.
5. Recover $\tilde{\mathbf{U}}_k = \mathbf{Q}\hat{\mathbf{U}}$ and the $k$ leading eigenvalues.

The dominant cost is the $2q+2$ matrix–matrix products with $\mathbf{G}$, each $O(n^2k)$, a substantial saving over $O(n^3)$ for $k \ll n$. A deterministic Krylov (Lanczos) method computes the same leading eigenpairs at comparable cost, and in our benchmarks gave the same accuracy with similar speed; RSVD needs a fixed number of passes over $\mathbf{G}$, which suits settings where $\mathbf{G}$ is only available through matrix products. The two-sided projection in step 4 is required: for a symmetric matrix, the one-sided form $\mathbf{B} = \mathbf{Q}^\top\mathbf{G}$ followed by an SVD identifies the correct subspace but not the eigenvectors within it, and silently returns a wrong low-rank $\mathbf{G}$.

**Rank selection criteria.** Each approximate solution $\hat{\mathbf{c}}_k$ was evaluated against the true $\mathbf{g}$ and $\mathbf{G}$, giving the gain difference, the signed coancestry difference and the exact objective gap $\delta_k$ relative to the full-dense solution (main text). Five rules for the recommended rank were compared, each taking the smallest tested rank that satisfies it on the replicate mean:

1. *Gain*: gain difference $< 1\%$ (the rule of earlier drafts).
2. *Objective 1%*: $\delta_k < 1\%$ (used in the main text).
3. *Objective 0.5%*: $\delta_k < 0.5\%$.
4. *Gain and coancestry*: gain difference $< 1\%$ and absolute coancestry difference $< 5\%$.
5. *Stabilisation*: the exact objective of $\hat{\mathbf{c}}_k$, computed from the selected candidates only, changes by less than 0.1% to the next tested rank. This rule needs no dense solve.

The gain rule consistently chose too low a rank: at the ranks it selected, coancestry exceeded the optimum by up to 51% (Norway spruce, $\gamma = 100$) and the objective gap reached 6% (Table S6). The stabilisation rule never chose a lower rank than the 1% objective rule, and did not settle within rank 100 wherever the 1% objective rule also failed or only just succeeded.

DRAFT

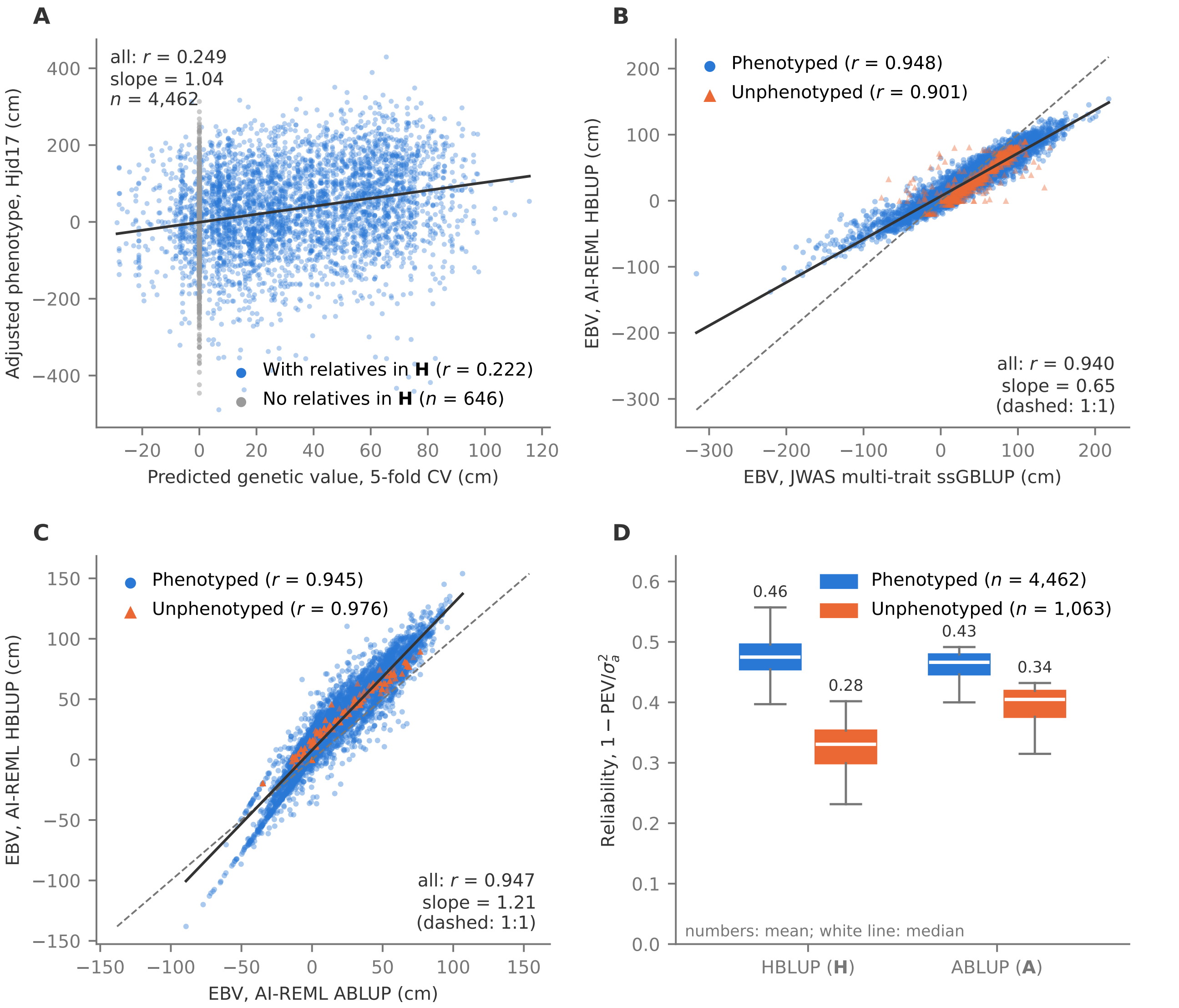


**Fig. S6. Single-step HBLUP AI-REML evaluation of Hjd17, Norway spruce.** **(A)** Five-fold cross-validated predicted genetic values versus phenotypes adjusted for trial and plot effects ($n = 4{,}462$); grey points are unrelated reference trees (unimproved material with genetic groups as parents), which have no relatives in $\mathbf{H}$ and therefore receive a zero cross-validated prediction. **(B)** HBLUP EBVs versus multi-trait JWAS posterior means. **(C)** HBLUP versus ABLUP EBVs. **(D)** Reliability $(1 - \mathrm{PEV}/\sigma_a^2)$ for phenotyped and unphenotyped trees under HBLUP and ABLUP. Solid lines: OLS fits; dashed lines: 1:1.

**Table S6.** Recommended rank under five rules

| Dataset | $\gamma$ | Gain | Obj. 1% | Obj. 0.5% | Gain + coan. | Stabilisation |
|---|---|---|---|---|---|---|
| Norway spruce | 100 | 5 | 10 | 15 | 10 | 25 |
| | 200 | 10 | 15 | 20 | 20 | 25 |
| | 400 | 10 | 25 | 30 | 25 | 30 |
| | 1000 | 25 | 30 | 30 | 30 | 50 |
| | 3000 | 30 | 30 | 40 | 30 | 40 |
| QTL-MAS 2010 | 0.1 | 5 | 5 | 15 | 30 | 5 |
| | 0.5 | 30 | 50 | 75 | 40 | 75 |
| | 1 | 30 | 100 | >100 | 75 | >100 |
| | 2 | 75 | >100 | >100 | >100 | >100 |
| | 5 | >100 | >100 | >100 | >100 | >100 |
| | 10 | >100 | >100 | >100 | >100 | >100 |

Tested ranks: 5, 10, 15, 20, 25, 30, 40, 50, 75, 100. >100: not reached within the tested ranks. Means over five RSVD replicates.

**Factorizing the uncertainty covariance in robust OCS.** Two source structures for $\boldsymbol{\Omega}$ behave differently under factorization. A posterior sample covariance from an MCMC analysis, $\hat{\boldsymbol{\Omega}} = \frac{1}{S-1}\mathbf{X}\mathbf{X}^\top$ for $S$ centred post-burn-in draws in $\mathbf{X} \in \mathbb{R}^{n \times S}$, is exactly rank $\leq S-1$, so $\mathbf{F}_\Omega = \mathbf{X}/\sqrt{S-1}$ is an exact factor with no approximation error and guaranteed positive semidefiniteness. A REML prediction error covariance matrix, $\mathbf{\Omega} = \sigma_a^2\mathbf{H} - \sigma_a^4\mathbf{H}_{\cdot o}\mathbf{P}\mathbf{H}_{o\cdot}$ with $o$ indexing phenotyped individuals and $\mathbf{P}$ the REML projection matrix, is strongly diagonally dominant and not low rank: for the Norway spruce dataset its 100 largest eigenvalues hold only 10% of its trace, and a pure rank-$r_\Omega$ factor approximates it poorly (its diagonal alone misstates $\sqrt{\mathbf{c}^\top\mathbf{\Omega}\mathbf{c}}$ by 28% for OCS-like contribution vectors).

We therefore use the same low-rank-plus-exact-diagonal form applied to $\mathbf{G}$,

$$\mathbf{\Omega} \approx \mathbf{F}_\Omega\mathbf{F}_\Omega^\top + \mathrm{diag}(\mathbf{d}_\Omega), \qquad \mathbf{d}_\Omega = \mathrm{diag}(\mathbf{\Omega}) - \mathrm{rowsums}(\mathbf{F}_\Omega \circ \mathbf{F}_\Omega),$$

with $\mathbf{F}_\Omega$ from a rank-$r_\Omega$ RSVD of $\mathbf{\Omega}$. The correction is non-negative because the residual is positive semidefinite, so the approximation is itself positive semidefinite — which matters, since $\mathbf{\Omega}$ sits under a square root in the robust objective — and both $\mathbf{\Omega}\mathbf{c}$ and $\sqrt{\mathbf{c}^\top\mathbf{\Omega}\mathbf{c}}$ cost $O(nr_\Omega)$ rather than $O(n^2)$. At $r_\Omega = 30$ the approximation reproduced the exact uncertainty term to within 0.3% for the OCS solution and 0.6% for the ROCS solution.

Embedding $\mathbf{G}$ and $\mathbf{\Omega}$ in a single block-diagonal matrix and decomposing once would give no benefit: the eigendecomposition decouples exactly into the two blocks, while forcing a shared rank budget across two spectra with different decay characteristics at higher memory cost.

### Supplementary Note 3: GBLUP analysis of the QTL-MAS 2010 dataset

The simulated pedigree of the 14th QTL-MAS workshop (2) comprises $n = 3{,}226$ individuals over five generations, founded by 5 males and 15 females. Two correlated traits were simulated under a model with additive, epistatic and parent-of-origin effects; only the continuous trait was analysed here. Phenotypes were available for the $n_{\text{pheno}} = 2{,}326$ individuals of the first four generations, while the $n_{\text{pred}} = 900$ individuals of the fifth generation had none and relied on genomic relatedness alone. Of the 10,031 simulated SNPs, 9,723 passed quality control (minor allele frequency $> 0.05$, call rate $> 0.95$).

GEBVs for all pedigree members were estimated by GBLUP fitted with AI-REML, with sex as a fixed effect and $\mathbf{G}$ constructed following VanRaden (3). Estimates, model fit criteria and prediction accuracies are given in Table S2, and predicted against observed values in Figure S2. All $n = 3{,}226$ pedigree members, phenotyped or not, were eligible candidates in the subsequent OCS, under the equal sex-contribution constraint $\sum_{i\in\mathcal{F}} c_i = \sum_{j\in\mathcal{M}} c_j = 50$ (contributions summing to 100).

DRAFT
